# Persistent synaptic inhibition of the subthalamic nucleus by high frequency stimulation

**DOI:** 10.1101/2021.10.20.465131

**Authors:** Leon A Steiner, Andrea A Kühn, Jörg RP Geiger, Henrik Alle, Milos R Popovic, Suneil K Kalia, Mojgan Hodaie, Andres M Lozano, William D Hutchison, Luka Milosevic

**Affiliations:** Krembil Brain Institute, University Health Network, Canada; Department of Neurology, Charité-Universitätsmedizin Berlin, Germany; Berlin Institute of Health (BIH), Germany; Institute of Neurophysiology, Charité-Universitätsmedizin Berlin, Germany; KITE Research Institute, University Health Network, Canada; Institute of Biomedical Engineering, University of Toronto, Canada; Department of Surgery, University of Toronto, Canada; Department of Physiology, University of Toronto, Canada

**Keywords:** deep brain stimulation, Parkinson’s disease, synaptic dynamics, synaptic depression, subthalamic nucleus, substantia nigra pars reticulata

## Abstract

**Background:** Deep brain stimulation (DBS) provides symptomatic relief in a growing number of neurological indications, but local synaptic dynamics in response to electrical stimulation that may relate to its mechanism of action have not been fully characterized.

**Objective:** The objectives of this study were to (1) study local synaptic dynamics during high frequency extracellular stimulation of the subthalamic nucleus (STN), and (2) compare STN synaptic dynamics with those of the neighboring substantia nigra pars reticulata (SNr).

**Methods:** Two microelectrodes were advanced into the STN and SNr of patients undergoing DBS surgery for PD. Neuronal firing and evoked field potentials (fEPs) were recorded with one microelectrode during stimulation from an adjacent microelectrode.

**Results:** Excitatory and inhibitory fEPs could be discerned within the STN and their amplitudes predicted bidirectional effects on neuronal firing (p = .007). There were no differences between STN and SNr inhibitory fEP dynamics at low stimulation frequencies (p > .999). However, inhibitory neuronal responses were sustained over time in STN during high frequency stimulation, but not SNr (p < .001) where depression of inhibitory input was coupled with a return of neuronal firing (p = .003).

**Interpretation:** Persistent inhibitory input to the STN suggests a local synaptic mechanism for the suppression of subthalamic firing during high frequency stimulation. Moreover, differences in the resiliency versus vulnerability of inhibitory inputs to the STN and SNr suggest a projection source- and frequency-specificity for this mechanism. The feasibility of targeting electrophysiologically-identified neural structures may provide insight into how DBS achieves frequency-specific modulation of neuronal projections.

## Introduction

Electrical high frequency stimulation (HFS) of the subthalamic nucleus (STN) has proven to be a reliable chronic intervention for the motor symptoms of Parkinson’s Disease (PD).^1^ Despite its therapeutic success, it is not fully understood how deep brain stimulation (DBS) interacts with neural tissue on a cellular level in humans. The rise of optogenetic manipulation in animal models of PD has established tools to investigate DBS mechanisms of action^2–4^ and engage with neuronal microcircuitries *in vivo* with unprecedented, millisecond precision.^5^ However, optogenetic control is not yet feasible in humans and fundamentally differs from DBS, which is delivered in the form of electrical impulses that non-selectively activate nearby neural elements. Intriguingly, recent translational advances have shown that knowledge of circuit architecture obtained through optogenetic experiments can be used to design electrical stimulation paradigms that achieve similar specificity in neurocircuit control which can yield substantial functional improvements in behavioral effects.^6^ To capitalize on the conceptual advances provided by optogenetic interrogation and to investigate the mechanism of action of DBS, we present an electrical stimulation framework that allows for the discernment of distinct neuronal projections and their effects on neuronal activity in the human basal ganglia.

In patients undergoing DBS surgery, synaptic readouts and corresponding alterations in single neuron firing in response to microstimulation have recently been reported to vary *across* brain regions according to the distributions of inhibitory and excitatory inputs.^7^ Analogous to recent findings in acute rodent brain slices,^8^ microstimulation may further allow the discernment of selective activation of excitatory and inhibitory projections *within* individual brain structures. Indeed, STN afferent inputs consist of excitatory “hyperdirect pathway” projections from the cortex and inhibitory “indirect pathway” projections from the globus pallidus externus (GPe). Stimulation of incoming projections has been implicated in the DBS mechanism of action and has been associated with a cessation of neuronal firing.^3^ Furthermore, direct optogenetic suppression of subthalamic somatic firing by ultrafast opsins has been shown to be therapeutic in the 6-hydroxydopamine rodent model of PD.^9^ It has also been shown that STN neuronal suppression could be achieved by high-frequency electrical microstimulation at settings of an equivalent electrical energy to those that are clinically effective.^10^ Despite these insights, the synaptic mechanism by which electrical stimulation achieves suppression of neuronal firing remains contentious. In patients with Parkinson’s disease, there is evidence that DBS can modulate inhibitory synaptic control in the basal ganglia,^11,12^ but evidence on plasticity of synaptic inhibition in the STN, the prime target of DBS, is missing.

Taking advantage of the unique opportunities of intraoperative microelectrode recordings in patients with Parkinson’s disease, the present study investigates how local extracellular microstimulation can be used for the discernment of incoming projections to the STN. Stimulation was delivered form one microelectrode while evoked fields and associated changes in neuronal activity were recorded from the adjacent microelectrode. This set-up has previously been used to suggest that extracellular electrical impulses could elicit excitatory or inhibitory changes to neuronal firing by activation of presynaptic terminals^7^ (although it is understood that other neural elements are also activated with stimulation pulses). Moreover, these presynaptic activations and changes to neuronal firing were also associated with synaptic readouts in the form of evoked field potentials in the context of neuronal inhibition.^12–14^ In this study, we (1) tested whether local extracellular microstimulation can selectively up- or downregulate neuronal activity in the human STN by recruitment of excitatory and inhibitory projections; (2) probed the robustness of inhibitory projections to the STN at low and high stimulation frequencies; and (3) compared inhibitory synaptic dynamics in STN, an established target of DBS in PD, to substantia nigra pars reticulata (SNr), a proposed DBS target for gait dysfunction.^15^ While HFS is routinely applied in the clinical application of STN-DBS, there is accumulating evidence to suggest that low frequency stimulation may be more beneficial in the SNr;^16,17^ however, there are currently no synaptic rationales to substantiate such clinical observations.

## Methods

### Patients and neurons

51 STN and 14 SNr neurons from 28 patients with PD were assessed in this study. All experiments conformed to the guidelines set by the Tri-Council Policy on Ethical Conduct for Research Involving Humans and were approved by the University Health Network Research Ethics Board and each patient provided written informed consent.

### Recording and stimulation

Brain recordings were acquired during awake DBS surgeries (OFF-medication) using two microelectrodes (600 μm apart, 0.1-0.4 MΩ impedances; Fig. 1) on a stainless-steel intracranial guidetube which shared a common ground. Recordings were obtained at ≥10kHz sampling frequency using two Guideline Systems GS3000 (Axon Instruments, Union City, USA) and digitized using a CED1401 data acquisition system (Cambridge Electronic Design). Microstimulation was delivered using a constant-current stimulator (Neuro-Amp1A, Axon Instruments, Union City, USA).

**Figure 1:**
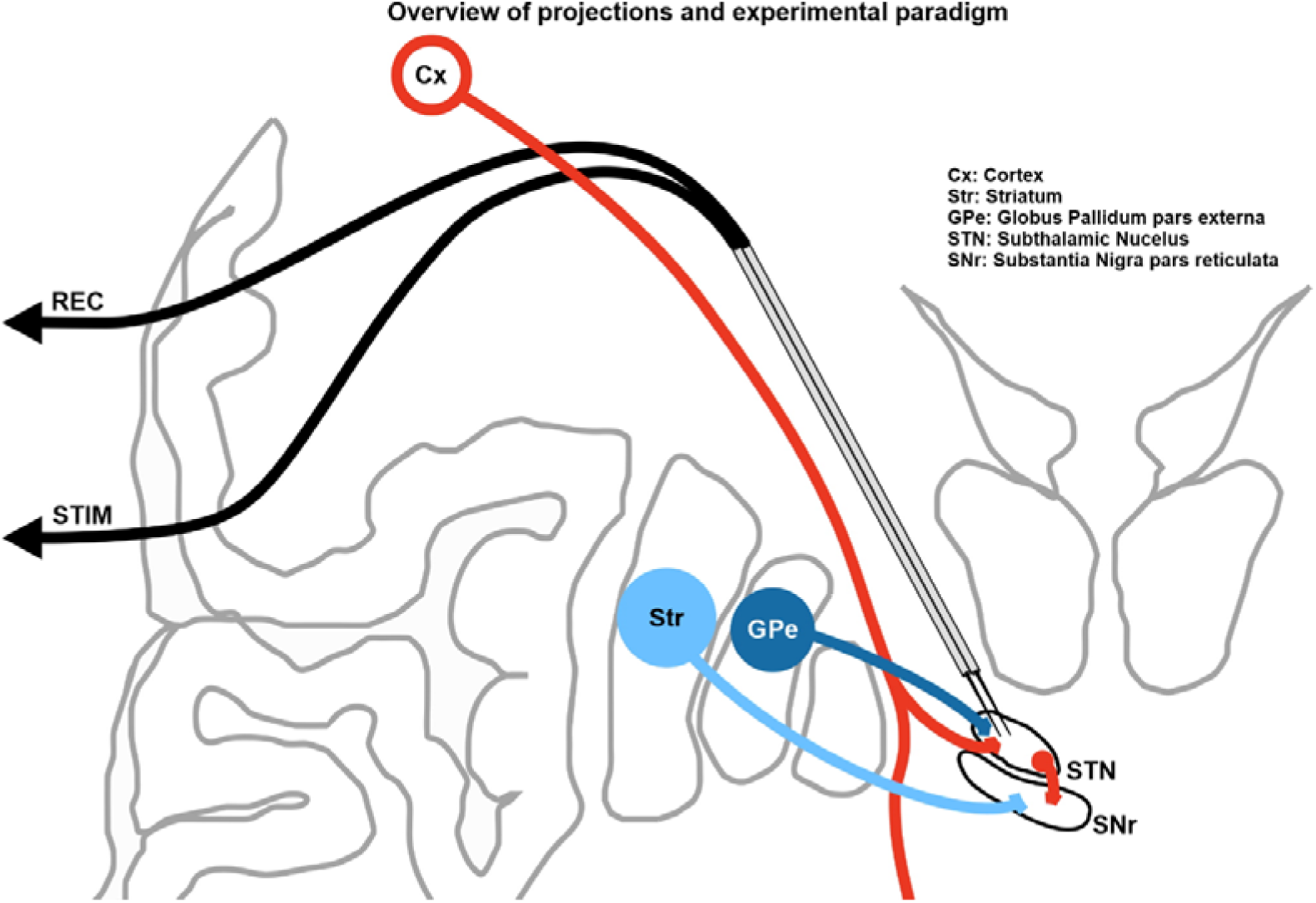
Overview of projections and experimental paradigm. Red line represents excitatory corticofugal neurons. Axon-collaterals of these neurons that project to the STN have been conceptualized as the so-called hyperdirect pathway. Dark blue line represents pallidal projection to the STN. Light blue line represents striatal projection to the SNr. Two closely spaced microelectrodes were advanced into the STN and SNr of patients undergoing DBS surgery for Parkinson’s disease. Neuronal activity was recorded with one microelectrode during stimulation trains from the adjacent microelectrode.

### Experimental protocols

Neuronal firing and evoked field potentials (fEPs) were recorded with one microelectrode during stimulation from an immediately adjacent microelectrode within the same structure. While recording from an STN or SNr neuron, stimulation trains of 100 µA at 0.3 ms biphasic pulses were applied at varied frequencies. To discern fEPs and changes in action potential firing in response to single stimuli in the STN, stimulation was applied at 1-5 Hz (10 s of each frequency) and pooled for subsequent analysis. To contrast synaptic inhibitory dynamics between STN and SNr at low and high stimulation frequencies, trains of 10 Hz and 100 Hz were applied for 5 s and 0.5 s, respectively (50 stimuli in each train). In a subsequent step, we investigated changes in action potential firing during longer duration trains (10 s) of 100 Hz stimulation, delivered to a subset of SNr neurons, as well as to a separate set of STN neurons. Please refer to Supplementary Table 1 for data summary and sources.

### Offline analyses and statistics

Statistical analyses were performed using Excel (Microsoft, WA, USA), SPSS Version 23.0 (IBM Corp, NY, USA), and Matlab 2018b (MathWorks Inc, MA, USA). Figures were created using the CorelDraw Graphics Suite (Corel, ON, Canada). Boxplots (central line, median; box, 25 % – 75 %; maximal whisker length, 1 time the interquartile range; data points beyond the whiskers displayed using “+”) were used to illustrate sample distributions. Stimulus-locked evoked field potentials (fEPs) were measured (peak minus pre-stimulus baseline) from the raw recordings. Synaptic fields were distinct from fiber volleys as illustrated in Fig. S1. To assess neuronal firing, data were high pass filtered (≥ 250 Hz) after artifact removal/blanking (required to avoid filter-related distortion), then template matching was done using a principal component analysis method in Spike2 (Cambridge Electronic Design, UK).

For Fig. 2, responses to single stimulation pulses at 1-5 Hz were averaged and analyzed taking into account both positive- and negative-going deflections 2-20 ms after the stimulation artifact. Furthermore, a composite measure (i.e. positive-minus negative-going deflection amplitudes, see Supplementary Fig. S2) was calculated. These stimulus-locked fEP amplitudes were normalized by the diving them by the average pre-stimulus rectified local field potential amplitude. This resulted in three data points for each neuron: the normalized positive-going component of the fEP, the normalized negative-going component of the fEP, and the normalized composite fEP, were each assigned to the same change in average firing rate, that was calculated as the change in the 50 ms post-stimulus interval in comparison to a 45 ms prestimulus baseline. The 5 ms bin in which the stimulus artifact occurred was not included in the analysis (gray bar in Fig. 2A/B/D). The stimulus-locked fEP amplitudes were then correlated (Pearson) with peristimulus changes in the firing rate. This was done for the positive-going, the negative-going and composite fEPs separately, and p-values were subsequently corrected for multiple comparisons using the Bonferonni correction (three comparisons). This established the inhibitory nature of positive-going fEPs (see Results), that were associated with brief pauses of somatic firing and are referred to as inhibitory fEPs in all further parts of the manuscript. For Fig. 3, stimulation frequency dependent dynamics of inhibitory evoked potentials in the STN were compared to inhibitory fEPs in the SNr. Neurons with discernable inhibitory fEPs at low and high stimulation frequencies were included in this analysis (Supplementary Table 1). Dynamics of inhibitory fEPs at low (LFS; 10 Hz) and high (HFS; 100 Hz) “frequency” (within-subject factor) stimulation were compared between “structures” (STN and SNr; between-subject factor), investigating the paired pulse (2^nd^/1^st^), as well as 5^th^/1^st^, 20^th^/1^st^, and 40^th^/1^st^ fEP “ratios” (within-subject factor) using a 2×2×4 split-plot repeated measures ANOVA design. For posthoc analysis, Bonferroni-corrected (four comparisons) two-tailed paired t-tests were used to compare effects between STN-LFS vs. STN-HFS, SNr-LFS vs. SNR-HFs, STN-LFS vs. SNr-LFS, and STN-HFS vs. SNr-HFS. For Fig. 4C/E, the relationship between the dynamics of the inhibitory fEPs and changes to firing rates over 10 s of stimulation at 100 Hz was investigated in both STN and SNr (2-tailed paired t-tests). To visualise changes of firing rate during ongoing stimulation over time, a plot of the average firing rate during the 10 s stimulation train was constructed for both STN and SNr neurons (Fig 4D). To discern differences between the suppression of firing at different time points statistically, serial Bonferroni-corrected (four comparisons) t-tests for four 2.5 s windows were conducted (Supplementary Fig. S3). For any neuron that displayed spiking during ongoing 100 Hz stimulation (n = 9 out of 24; 37.5 % for STN, n = 6 out of 7; 85,7 % for SNr), the exact timing of spiking within the 10 ms interval between stimuli was analyzed in Fig. 4G/H. For group level statistics, the 10 ms interstimulus interval was divided into two 5 ms windows. Separate t-tests were performed to compare the firing rate in the first 5ms second window against the firing rate in the second 5 ms second in both STN and SNr, respectively.

**Figure 2:**
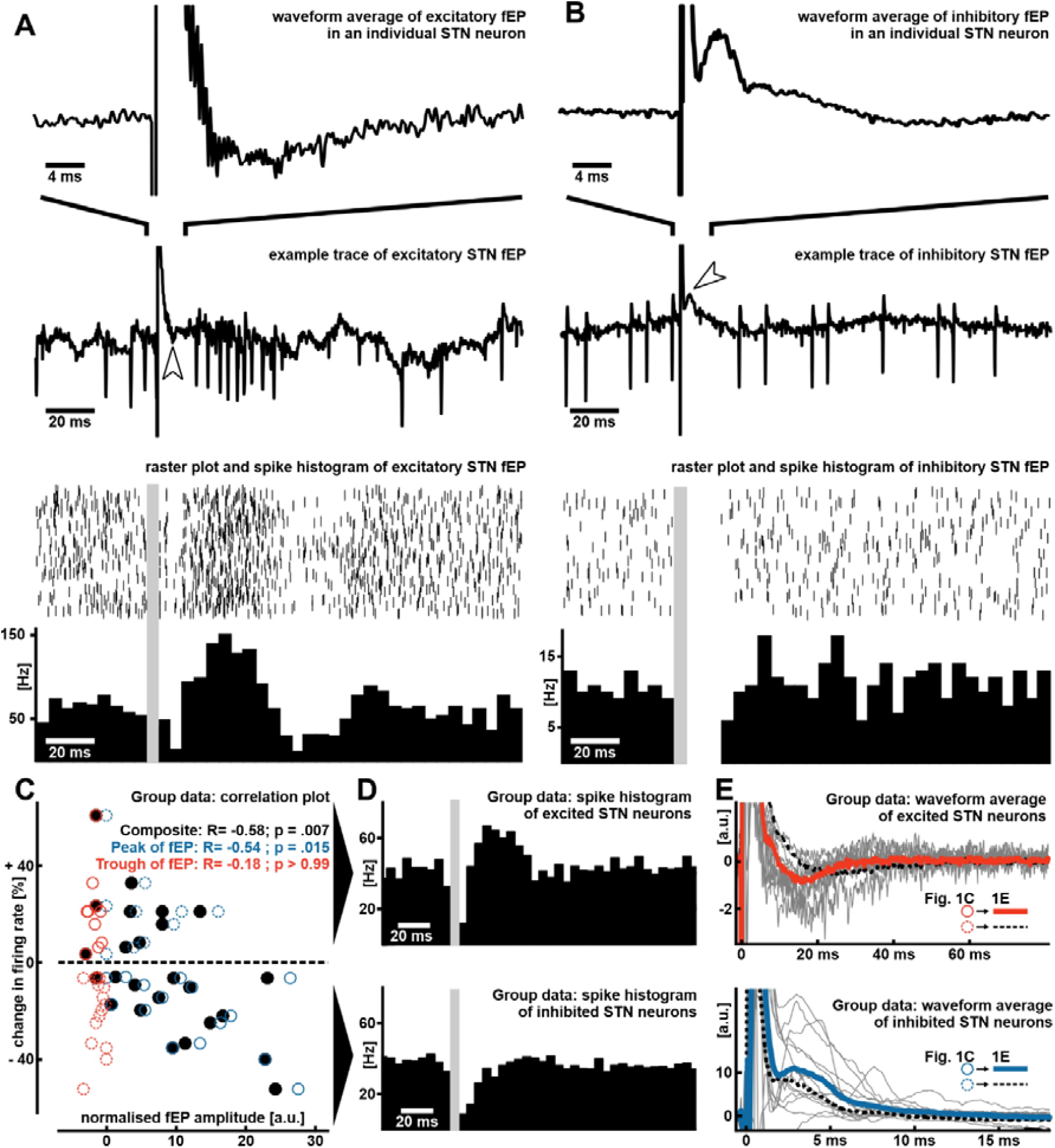
fEPs are linked to the change of firing rate of human STN neurons. (A&B) Stimulus-locked evoked potentials and action potential firing in excited and inhibited example STN neurons. Upper panels: Averaged negative and positive going evoked potentials at high temporal resolution. Middle panels: Raw traces of evoked potentials and firing rate changes in response to single stimuli. Arrowheads indicate evoked potentials. Lower panels: Raster plots and spike histograms of the same example neurons. (C) Group data show the relationship between LFP-normalized fEP amplitudes and firing rate changes in the 50 ms poststimulus time window compared to prestimulus baseline across neurons (n = 25 neurons from 14 patients). Note that each neuron is represented by three data points, one of each color, representing the positive going (blue), negative going (red) and composite (black) fEP (see Supplementary Figure S2 for illustration how the composite evoked potential was derived). The dashed line at zero separates excited end inhibited neurons. R- and p-values taken from multiple comparison corrected Pearson correlations. Dashed datapoints indicate negative going fEPs in inhibited neurons and positive going fEPs in excited neurons. (D) Upper panel: Average spike histogram of excited neurons (n = 10). Lower panel: Average spike histogram of inhibited neurons (n = 15). (E) Upper panel: Grey – individual fEP waveforms of excited neurons (n = 10), Red – Average fEP waveform of excited neurons, dotted black – average fEP of inhibited neurons for comparison. Lower panel: Grey – individual fEP waveforms of inhibited neurons (n = 15), Blue – average fEP waveform of inhibited neurons, dotted black – average fEP waveform of excited neurons for comparison.

**Figure 3:**
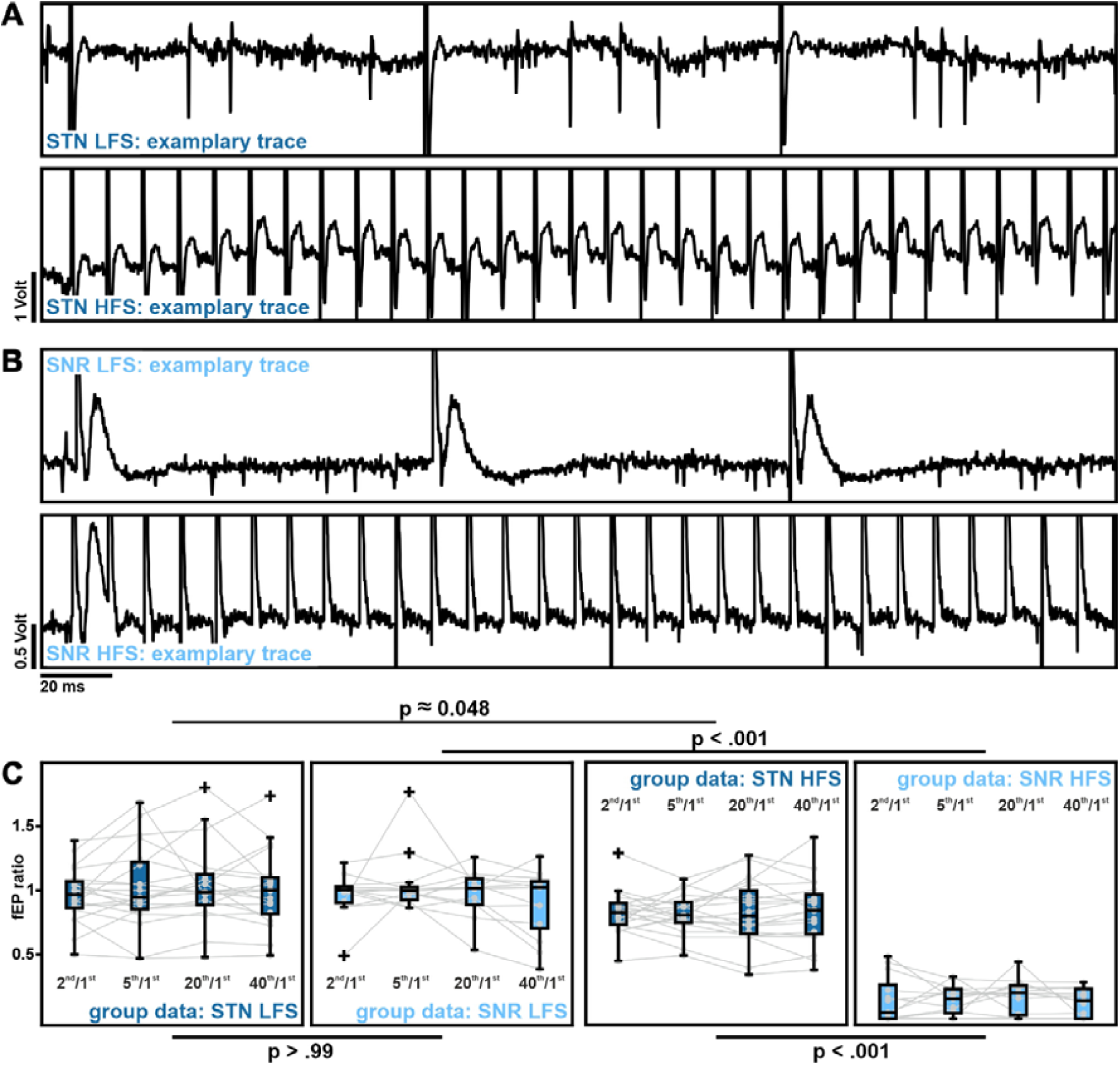
Inhibitory evoked potentials are sustained at HFS in STN but not SNr. (A) Example traces displaying inhibitory fEPS at low (10 Hz; upper panel) and high frequency stimulation (100 Hz; lower panel) in the STN. (B) Example traces displaying inhibitory fEPS at low (10Hz; upper panel) and high frequency stimulation (100 Hz; lower panel) in the SNr. (C) Group data show distribution of fEP ratios (second/first, fifth/first, twentieth/first, fortieth/first) for STN LFS, SNR LFS, STN HFS, and SNR HFS (n_STN_ = 20 neurons from 11 patients, n_SNr_ = 13 neurons from 9 patients). Grey data points represent individual cells. P-values are taken from multiple comparison corrected posthoc t-tests.

**Figure 4:**
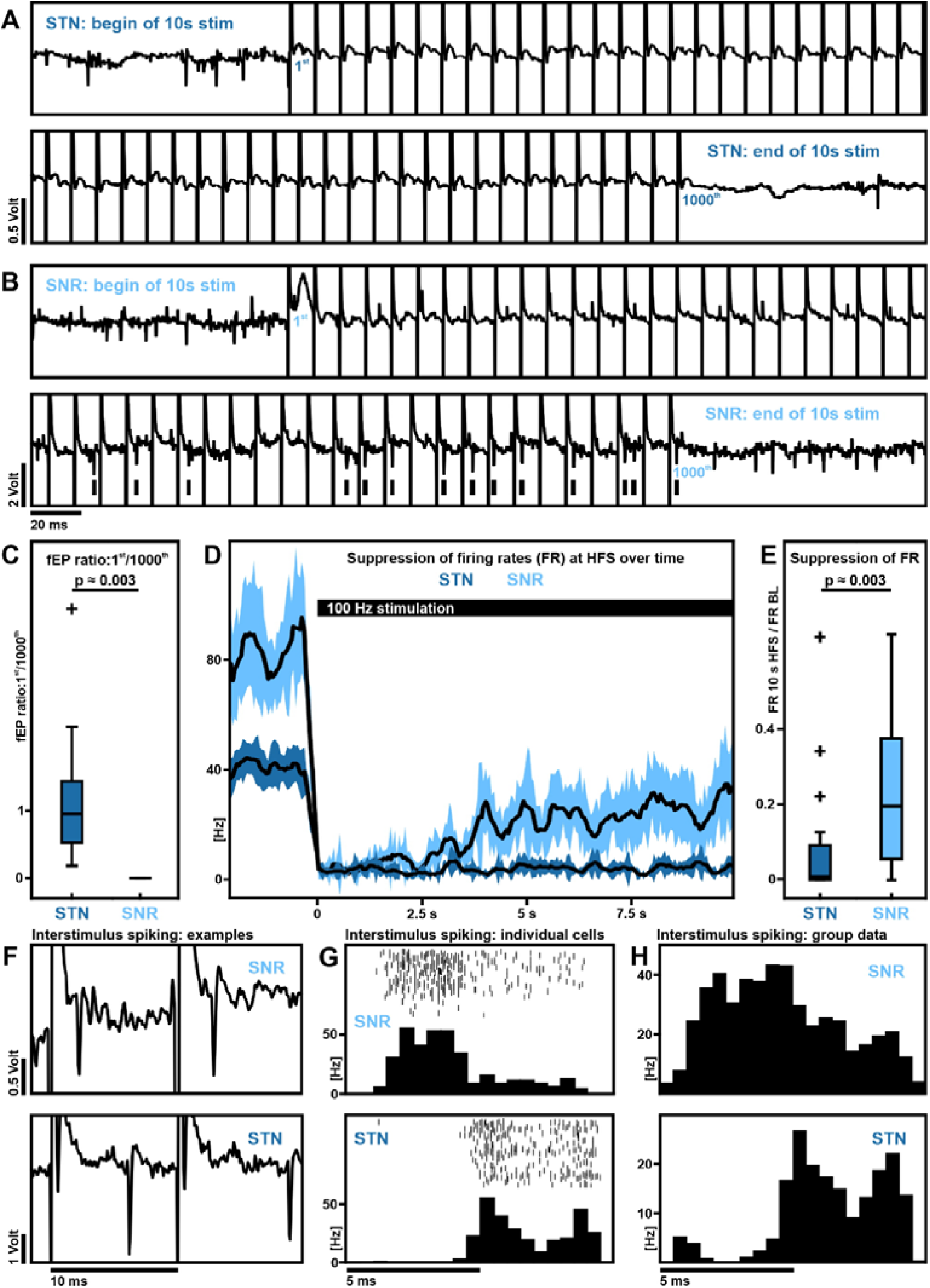
Continuous suppression of firing rates at HFS in STN but not SNr. (A) Example traces of the beginning (upper panel) and the end (lower panel) of 10 s stimulation interval at HFS in the STN featuring sustained inhibitory evoked potentials and continued suppression of action potential firing. (B) Example traces of beginning (upper panel) and end (lower panel) of 10s stimulation interval at HFS in the SNr featuring depressed inhibitory evoked potentials and a re-occurrence of action potential firing. Vertical lines indicate action potentials during ongoing stimulation. (C) Group data show distribution of fEP ratios (thousandth/first) for STN and SNr (n_STN_ = 19 neurons from 10 patients, n_SNr_ = 7 neurons from 7 patients). (D) Group data show dynamics of average firing rate for STN and SNR neurons over the 10s stimulation period (n_STN_ = 24 neurons from 10 patients, n_SNr_ = 7 neurons from 7 patients). Shadow indicates standard error. (E) Group data show distribution of average firing rates of the entire 10 s stimulation window (n_STN_ = 24 neurons from 10 patients, n_SNr_ = 7 neurons from 7 patients). (F) Example traces of interstimulus spiking for SNr (upper panel) and STN (lower panel). (G) Raster plots and spike histograms of interstimulus spiking for the same example neurons shown in (F). (H) Upper panel: Average histogram of interstimulus spiking for SNr neurons (n_SNr_ = 6 neurons from 5 patients). Lower panel: Average histogram of interstimulus spiking for STN neurons (n_STN_ = 9 neurons from 4 patients).

## Data availability

Data are available at https://www.biorxiv.org/content/10.1101/2020.11.30.404269v1.

## Results

### Evoked field responses were linked to bidirectional changes in STN neuronal firing rates

Responses to single stimulation pulses showed that negative-going deflections in the fEPs were associated with a net increase of action potential firing the 50 ms after the stimulus (Fig. 2A), while positive-going deflections resulted in a pause of firing (Fig. 2B). Most stimulation sites yielded fEPs with both positive- and negative-going components. A composite fEP integrating these features was correlated with relative change in firing rate from the pre-stimulus baseline (Fig. 2C; Rho = −0.5812; p = .007). A larger composite fEP (found at 20/24 stimulation sites; dominated by an upward component) was associated with a decrease in the post-stimulus firing rate, suggesting an inhibitory effect of the upward deflection of the evoked response (henceforth referred to as an inhibitory fEP); while a smaller or net negative composite fEP was associated with an increase in the post-stimulus firing rate, indicating an excitatory effect of the downward deflection (henceforth referred to as an excitatory fEP). The amplitude of the inhibitory fEP correlated with the change in firing rate (Rho = −0.5422; p = .015), while the excitatory fEP did not (Rho = −0.1835; p > .999). To visualize fEPs on the group level, cells were grouped according to observed firing rate changes (Fig. 2D/E). This analysis provides evidence that electrical stimuli elicited selective increases or decreases in the firing rate of human STN neurons, and that evoked potential amplitudes (particularly composite and inhibitory fEPs) represent a viable substrate to predict these responses.

### Inhibitory evoked potentials were sustained with HFS in STN but not SNr

ANOVA analyses (Fig. 3) revealed stronger depression of inhibitory fEPs at HFS compared to LFS (main effect of frequency F_1,16.098_ = 157.647, p < .001) and stronger depression for SNr compared to STN (main effect of structure F_1,8.051_ = 41.305, p < .001). The main effect of ratio (2^nd^/1^st^, 5^th^/1^st^, 20^th^/1^st^ and 40^th^/1^st^ fEP amplitude) was not significant (F_3,0.083_ = 1.474, p = .227). Further, we observed a frequency-structure interaction (F_1,6.468_=63.341, p < .001), but not a ratio-structure (F_1,0.043_ = 2.376, p = .133), ratio-frequency (F_1,0.004_ = 0.153, p = .698), nor ratio-structure-frequency (F_1,0.025_ = 0.893, p = .352) interaction. For post-hoc analyses, ratios were averaged for each structure at a given frequency. Much stronger depression of fEPs was observed in SNr compared to STN at HFS (p < .001); but no difference between STN and SNr was found for ratios at LFS (p > .999). Within structures, there was a small difference between LFS and HFS ratios for STN (p = .048) and a large difference for SNr (p < .001). These results show that inhibitory fEPs in the STN are largely sustained at HFS, compared to rapidly and continuously depressed inhibitory fEPs in the SNr. At LFS, both STN and SNr show no difference in inhibitory fEP dynamics and there is no depression throughout the stimulation train in either structure.

### Continuous suppression of firing rates with HFS in STN but not SNr

Inhibitory fEPs were sustained during 10 s of stimulation at 100 Hz in STN, compared to fEPs in SNr which were rapidly and continuously suppressed during 10s of stimulation (Fig. 3A/B/C; p = .003). Further, the average firing rate was higher in SNr compared to STN neurons across the entire 10 s stimulation train (Fig. 4E; p = .003). To visualize the return of firing over time, a plot of the average firing rate during the 10 s stimulation was constructed for both STN and SNr neurons (Fig. 4D). The 10 s of stimulation were devided into four 2.5 s windows for further analysis (Supplementary Fig. S3). There was no difference for the first 2.5 s window (1^st^ window: mean_STN_ = 3.4, mean_SNr_ = 5.429, p > .999), but suppression of STN neurons was stronger than suppression of SNr neurons for the other three 2.5 s windows (2^nd^ window: mean_STN_ = 3.5, mean_SNr_ = 15.2, p = .015; 3^rd^ window: mean_STN_ = 4.3, mean_SNr_ = 22.51, p = .004; 4^th^ window: mean_STN_ = 4.1, mean_SNr_ = 26.8, p = .006). In a subset of STN neurons (n = 9 out of 24; 37.5 %), included in the analysis above, incomplete suppression of firing was observed. The timing of interstimulus spiking in this subset of STN neurons was compared to the timing of SNr neurons that displayed a return of firing (n = 6 out of 7; 85.7 %). In STN neurons, spiking was abolished almost completely in the first 5 ms window of the 10 ms interstimulus interval and, when present, occurred in the second 5 ms window (Fig. 4G/H; p = .008). For SNr, this relationship was not present, though spiking appeared to favour the first 5 ms window (p = .107). These data demonstrate that inhibitory fEPs were sustained over 10 s coupled with a continued suppression of firing rates for STN neurons. In contrast, the depression of SNr inhibitory fEPs was associated with a relative return of neuronal firing. In the subset of STN neurons that displayed incomplete suppression of spiking during ongoing stimulation, spiking robustly favoured the second half of the 10 ms interstimulus interval.

## Discussion

This study examined the frequency-specific mechanisms underlying synaptic control of basal ganglia structures using extracellular electrical stimulation. The experimental paradigm and analysis scheme used represent tools to discriminate inhibitory and excitatory afferent input to the STN in humans. Further, the results highlight the robustness of inhibitory projections onto STN neurons as a distinctive property that allows for continuously suppressed neuronal activity at high stimulation frequencies that are similar to those used in a clinical setting. In contrast, inhibitory input to the SNr depressed rapidly at HFS and the sustained depression was coupled with a relative return of SNr spiking. Ultimately, the structure-to-structure differences in synaptic dynamics may pave the way for electrophysiology-guided projection-specific interventions in humans and can provide synaptic rationales to guide future DBS programming.

### Discriminating inhibitory from excitatory synaptic fields in the human STN to identify recruited projections

While inhibitory SNr fEPs had previously been described in PD patients,^11^ STN fEPs are described here for the first time. SNr fEPs are robust and of large amplitude, in comparison to STN fEPs that are of comparatively lower amplitudes and far more heterogenous in terms of polarity (i.e. positive, negative, or a mix of both). This heterogenous presentation of STN fEPs nevertheless had a clear, bidirectional impact upon neuronal firing. We show that the inhibitory component of the fEP can explain the variability of the post-stimulus firing rate changes, while the excitatory components could not. The latter might be due to the fact that excitatory components in the fEP are partially occluded by the inhibitory component, as the peaks of postsynaptic currents in rodent STN slices have been shown to temporally overlap.^8^ Of note, the composite signal that integrates both inhibitory and excitatory components was best able to explain the variability of firing rate changes (Fig. 2C).

In contrast to excitatory inputs to the STN that are a composite of glutamatergic projections of variable origins, the vast majority of inhibitory GABAergic inputs to the STN come from the GPe.^18^ Thus, activation of GABAergic input to the STN can be expected to engage a specific neuronal circuit that has been suggested to orchestrate pathological basal ganglia activity.^19^ In its circuit-specificity, stimulation of inhibitory afferents to the STN may thus be comparable to optogenetic activation of subgroups of GPe neurons that project to the STN. In rodents, this circuit-intervention has been shown to achieve long-lasting motor recovery.^2,6^ Hence, the identification of inhibitory input to the STN in humans may hold translational potential for future studies selectively targeting this distinct projection.

### Frequency-dependent modulation of inhibitory projections to DBS targets

Recent computational studies have proposed that HFS leads to neuronal suppression by depression of synaptic currents.^7,20^ In this work, we demonstrate more nuanced site-specific synaptic mechanisms which underlie the control of somatic firing during HFS. Herein, we demonstrate that suppression of neuronal firing in STN is achieved by sustained inhibitory synaptic transmission during HFS; whereas inhibitory inputs to the SNr were subject to synaptic depression. These findings suggest a difference in the resiliency (versus vulnerability) to synaptic depression depending on the projection-specific source of the fibers activated (Fig. 5A). The reason for this difference may be ontogenetic: striatal neurons (which comprise the majority of inhibitory inputs to SNr)^21^ physiologically fire at low frequencies (5-10Hz) and are therefore likely more labile, while GPe firing rates are generally higher (40-60Hz),^22^ thus requiring an adapted synaptic organization to achieve reliable transmission. Further, while GPe neuronal firing is purportedly lower in Parkinson’s disease, it has been suggested that the strength of inhibitory GPe-subthalamic synaptic transmission is enhanced.^23^ Given the resilience of pallidal synaptic inhibition to the STN reported in this study, it is thus possible that DBS leverages this enhanced synaptic efficacy to promote maintained inhibitory drive at a high frequency. This is consistent with previous work in rodent brain slices that has indicated that GABAergic input to the STN is partially sustained at HFS^8^ and may also explain the therapeutic action of muscimol (GABA-A-agonist) microinjections into the STN that have been shown to reverse parkinsonian symptoms.^24,25^

**Figure 5:**
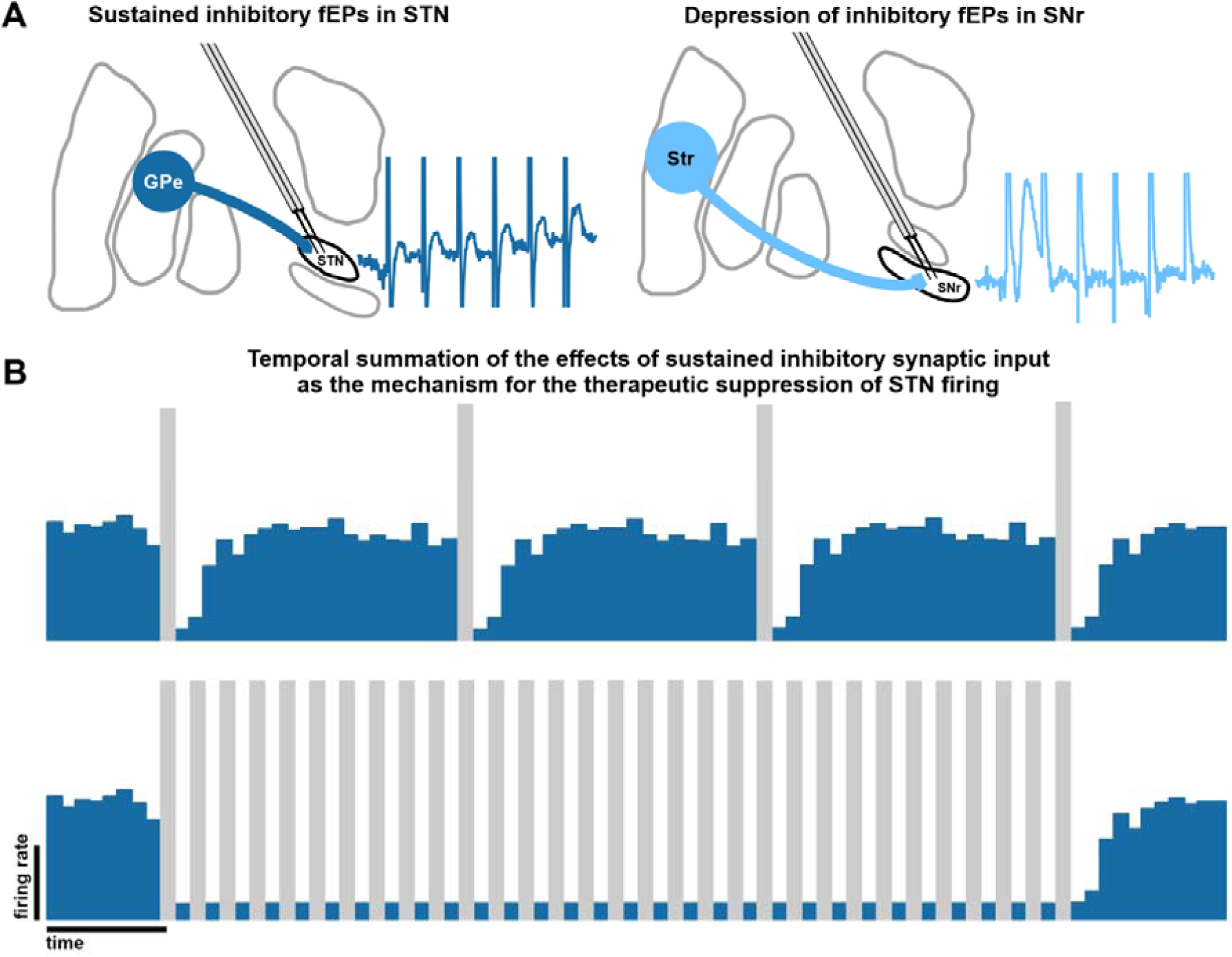
A local synaptic mechanism of DBS at the mercy of synaptic physiology. (A) Although small in amplitude, inhibitory fEPs were sustained at HFS in contrast to depression of inhibitory fEPs in SNr, indicative of a short-term depression synaptic mechanism. (B) Schematic: changes of firing rate in STN neurons. Stimulation pulses suppress STN firing at LFS for only short periods of time (upper panel) but if applied at high frequency these short periods summate to sustained suppression of STN firing (lower panel), likely representing a therapeutic mechanism of STN-DBS.

Despite somatic inhibition, STN efferent axons may nevertheless be entrained; which forms the basis of the so-called soma-axon-decoupling theory.^26^ To this end, evoked resonant neural activity^27^ has recently been hypothesized to be a substrate of STN efferent-driven recurrent inhibition from GPe.^28,29^ This interpretation would also be consistent with persistent pallidal synaptic inhibition at high frequencies described in this study, given that evoked resonant neural activity is a resilient signal which occurs at frequencies above 250 Hz.

In SNr, where strong and robust synaptic depression of inhibitory fEPs is achieved, there is a return of neuronal firing. This may be mediated by spontaneous ionic mechanisms such as sodium leak channels^30^ which operate independently from afferent synaptic inputs and may govern neuronal firing once tonic inhibition is lost due to the vulnerability of inhibitory inputs during HFS. Alternatively, presynaptic neuromodulation may be at play: GABA-B-dependent depression of subthalamo-nigral afferents that has been described in rodent brain slices^31^ has been shown to desensitize in the timescale of seconds in other preparations, potentially unleashing STN-SNr glutamatergic inputs in a time frame that may explain the temporal delay of firing in the SNr observed in this study. Future experiments assessing the effects of STN stimulation on SNr activity can help elucidate the synaptic dynamics of STN efferents and assess the importance of efferent vs afferent activations in the mechanisms of action of DBS.

### Towards synaptic dynamics informed deep brain stimulation

While the amplitude of inhibitory input and the corresponding pause in firing may be lesser in the STN compared to the SNr, the robustness of synaptic inhibitory input at HFS is specific to STN. Temporal summation of the inhibitory effect of these inputs at high frequencies may thus achieve a sustained suppression of pathologically overactive firing,^35^ potentially representing a synaptic rationale as to why HFS is clinically effective in STN (Fig. 5B), but not SNr.

In a subset of STN neurons we observed incomplete suppression of firing during trains of 100 Hz stimulation. Interestingly, firing in STN neurons occurred almost exclusively in the second 5 ms window of the 10 ms interstimulus interval, likely due to a weak, but sustained inhibitory drive. This relationship was not present in SNr, where the inhibitory drive was completely depressed and firing during 100 Hz stimulation tended to favor the first half of the 10 ms interstimuli interval. Because somatic firing is almost completely absent for the first 5 ms in STN, stimulation frequencies of above 100 Hz, where the interval between stimuli is less than 10 ms, would suppress somatic firing in STN to even higher degrees (under the assumption that GPe could follow these frequencies; not assessed here). This corresponds to clinical observations that have found that stimulation frequencies above 100 Hz or even the conventional 130 Hz can provide additional benefit in refractory or tremor dominant cases treated with STN-DBS.^32–34^

### Considerations and limitations

This study has provided synaptic evidence as to why recruitment of afferent inhibitory projections at different frequencies may differ in its success in achieving neuronal suppression. However, concerning the DBS mechanism of action, antidromic recruitment of afferents as well as recruitment of efferent axons by electrical stimulation must also be considered^36^ as neuronal firing at downstream targets of the stimulated structure has been shown to be regularized rather than supressed by subthalamic DBS.^37^ Interestingly, neurons that displayed incomplete suppression in our sample also showed highly patterned firing (Fig 4 G/H). Furthermore, DBS has been shown to suppress hypersynchronized oscillatory in the beta frequency band at both DBS target structures^38^ and synaptically coupled structures of the cortex.^39^ Despite this link between oscillatory and disease states, reflecting the importance of both distant and local effects of DBS, functional impairment can be controlled solely by suppression of somatic burst-firing patterns in the STN without the necessity to interfere with aggregate level oscillations.^40^ Ultimately, abolishing pathological patterns of firing might be comparable to a reversible functional lesion, a concept that has historically been suggested as crucial to the DBS mechanism of action.^41^

A notable limitation of the present report is that stimulation was applied for only short durations and the presented changes to synaptic dynamics should be validated over longer periods of time. Moreover, this study focused on neurophysiological effects of stimulation, but clinical or behavioural correlates were not quantified or correlated with these surrogates directly. Finally, there are differences in current density between the microstimulation applied here and macrostimulation that is used in the clinical application of DBS. However, as shown previously, the stimulation parameters used here (in particular, the 100 Hz microstimulation trains) are comparable in terms of total electrical energy delivered during clinically applied DBS macrostimulation.^10^ This same microstimulation at 100 Hz and 200 Hz applied to the ventral intermediate nucleus (Vim) of the thalamus was also effective at suppressing tremor.^42^

## Conclusion

In conclusion, this study establishes a framework to distinguish excitatory and inhibitory projections in the human STN and reveals frequency- and projection-dependent differences in inhibitory synaptic dynamics in response to extracellular electrical stimulation as is delivered in DBS therapy. In order to tune the target microcircuit in a functionally dependent manner, future stimulation paradigms may be critically informed by these dynamics, especially when exploring DBS in new targets and indications.

## Acknowledgements

The authors would like to thank the patients for their participation.

## Author contributions

Conceptualization: LAS, AAK, WDH, LM

Data curation: LAS, LM

Formal Analysis: LAS, LM

Funding acquisition: LAS, AAK, MRP, WDH, LM

Investigation: LAS, SKK, MH, AML, WDH, LM

Methodology: WDH, LM

Project administration: LAS, AAK, WDH, LM

Resources: MRP, SKK, MH, AML, WDH, LM

Software: LAS, LM

Supervision: AAK, LM

Validation: WDH, LM

Visualization: LAS, LM

Writing – original draft: LAS

Writing – review & editing: LAS, AAK, JRPG, HA, MRP, SKK, MH, AML, WDH, LM

