## Supplement for "Persistent synaptic inhibition of the subthalamic nucleus by high frequency stimulation"

**Supplementary material: Steiner et al. 2021**

***Supplementary Table 1: Experimental data summary.***

|  | **INCLUDED** | **EXCLUDED** (rationale to exclude) |
| --- | --- | --- |
| Fig. 1 | **Examples:**  Fig. 1A: STN19  Fig. 1B: STN12  **Group data:**  STN1-STN22, STN25-STN27 | **Group data:**  STN23 (amplitude not discernable due to signal distortion); STN24 (putative antidromic spike leading to waveform distortion) |
| Fig. 2 | **Examples:**  Fig. 2A: STN17  Fig. 2B: SNR9  **STN Group data:**  STN1-12, STN14, STN16-STN17, STN20-STN22, STN25-STN26  **SNR Group data:**  SNR1-SNR13 | **STN Group data:**  STN23 and STN24 (see above); STN13, STN15, STN 18, STN19 and STN27 (no clearly discernable inhibitory fEP at stimulation onset)  **SNR Group data:**  SNR14 (no inhibitory fEP discernable) |
| Fig. 3 | **Examples:**  Fig. 3A: STN67  Fig. 3B: SNR11  Fig. 3F,G (upper panels): SNR11  Fig. 3F,G (lower panels): STN50  **STN Group data (Fig. 3C):**  STN48-STN51, STN53-STN54, STN56-STN58, STN60-STN62, STN64, STN66-STN71  **STN Group data (Fig. 3D,E):**  STN48-STN71  **SNR Group data (Fig. 3C,D,E):**  SNR1, SNR3, SNR5, SNR7, SNR10, SNR11, SNR13  **STN Group data (Fig. 3H, lower panel):**  STN48, STN49, STN50, STN55, STN56, STN57, STN66, STN69, STN70  **SNR Group data (Fig. 3H, upper panel):**  SNR3, SNR5, SNR7, SNR10, SNR11, SNR13 | **STN Group data (Fig. 3C):**  STN52, STN55 (no clearly discernable inhibitory fEP at stimulation onset), STN59 (amplitude not discernable due to overlay of stimulation artifact), STN65 (no clearly discernable inhibitory fEP at stimulation onset)  **STN Group data (Fig. 3D,E):**  None excluded  **SNR Group data (Fig. 3C,D,E):**  SNR4, SNR6 and SNR12 (poor signal quality); SNR2, SN8, SNR9, SNR14 (no long recording available) |

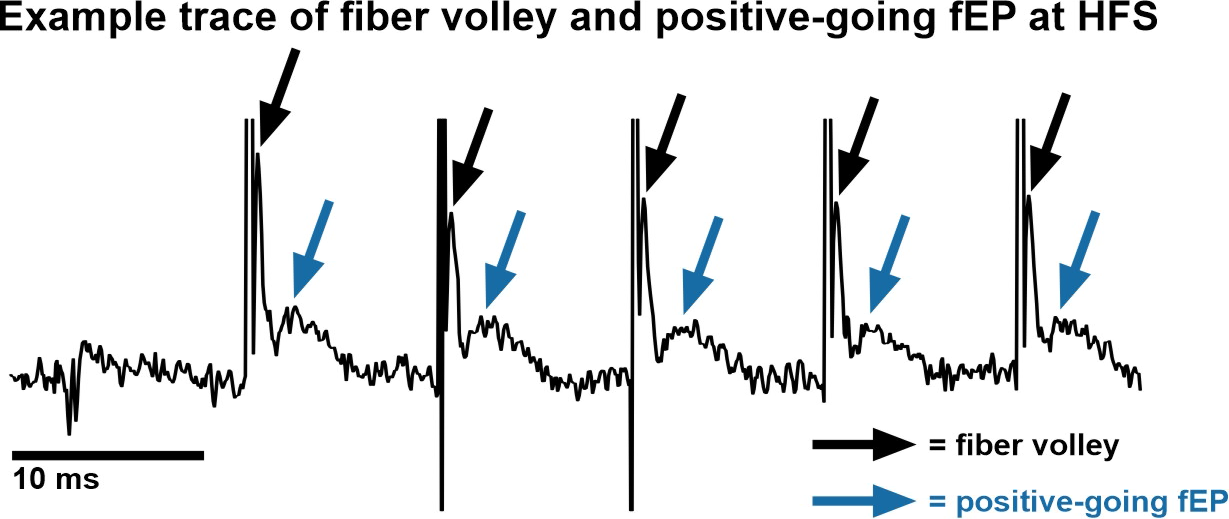

***Supplementary Figure S1: Example trace of fiber volley and positive-going fEPs at HFS.*** This example trace at high temporal resolution demonstrates that positive-going fEPs occurred as distinguishable signals without overlap with the fiber volley.

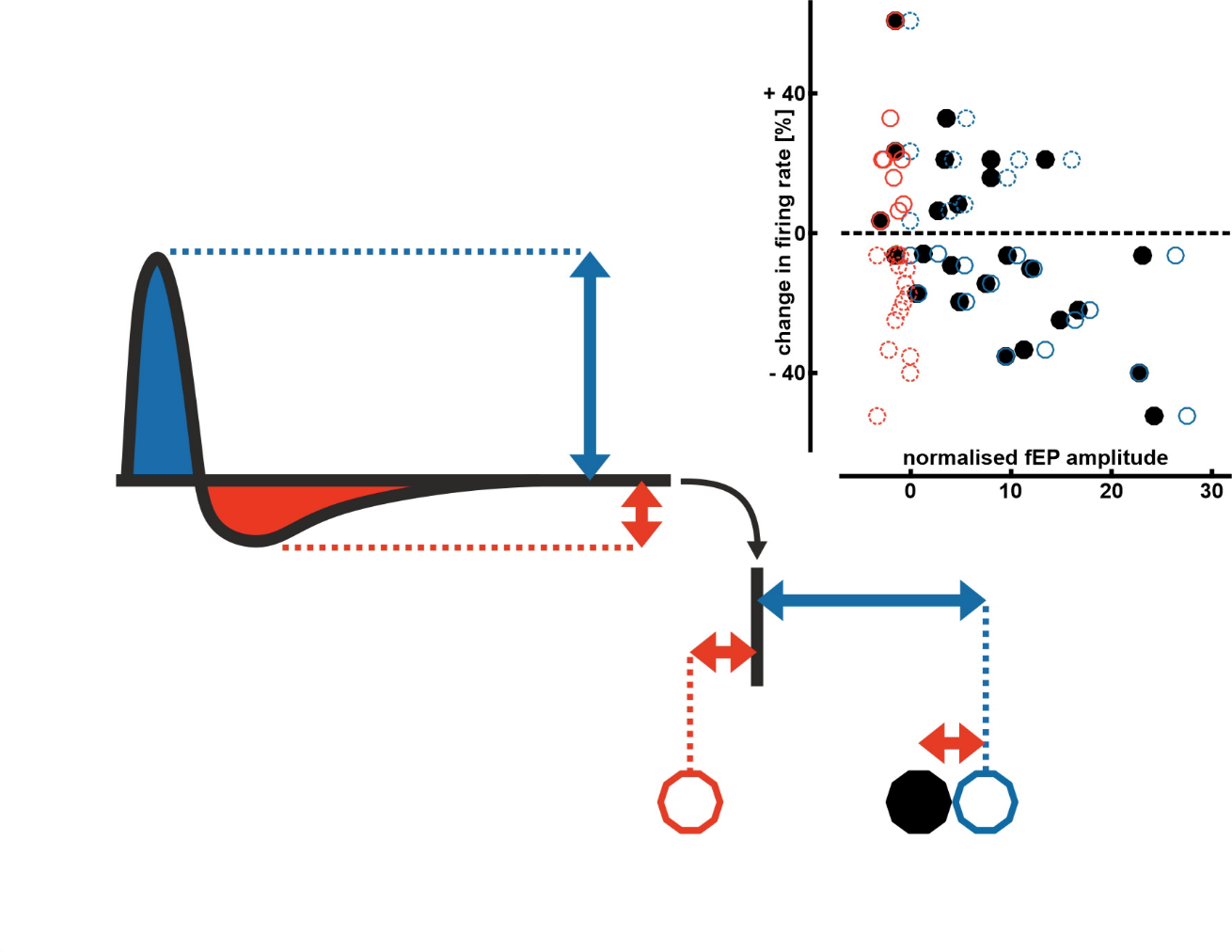
***Supplementary Figure S2: Schematic to illustrate how the composite evoked potential was derived from positive and negative going components of each signal.*** A composite fEP was calculated by subtracting the amplitude of the through of each fEP from the amplitude of its peak. This resulted in three data points for each neuron: positive-going component of the fEP (blue), negative-going component fEP (red) and composite fEP (black), each assigned to the same change in firing rate. Top right inset is taken from Figure 2C for reference.

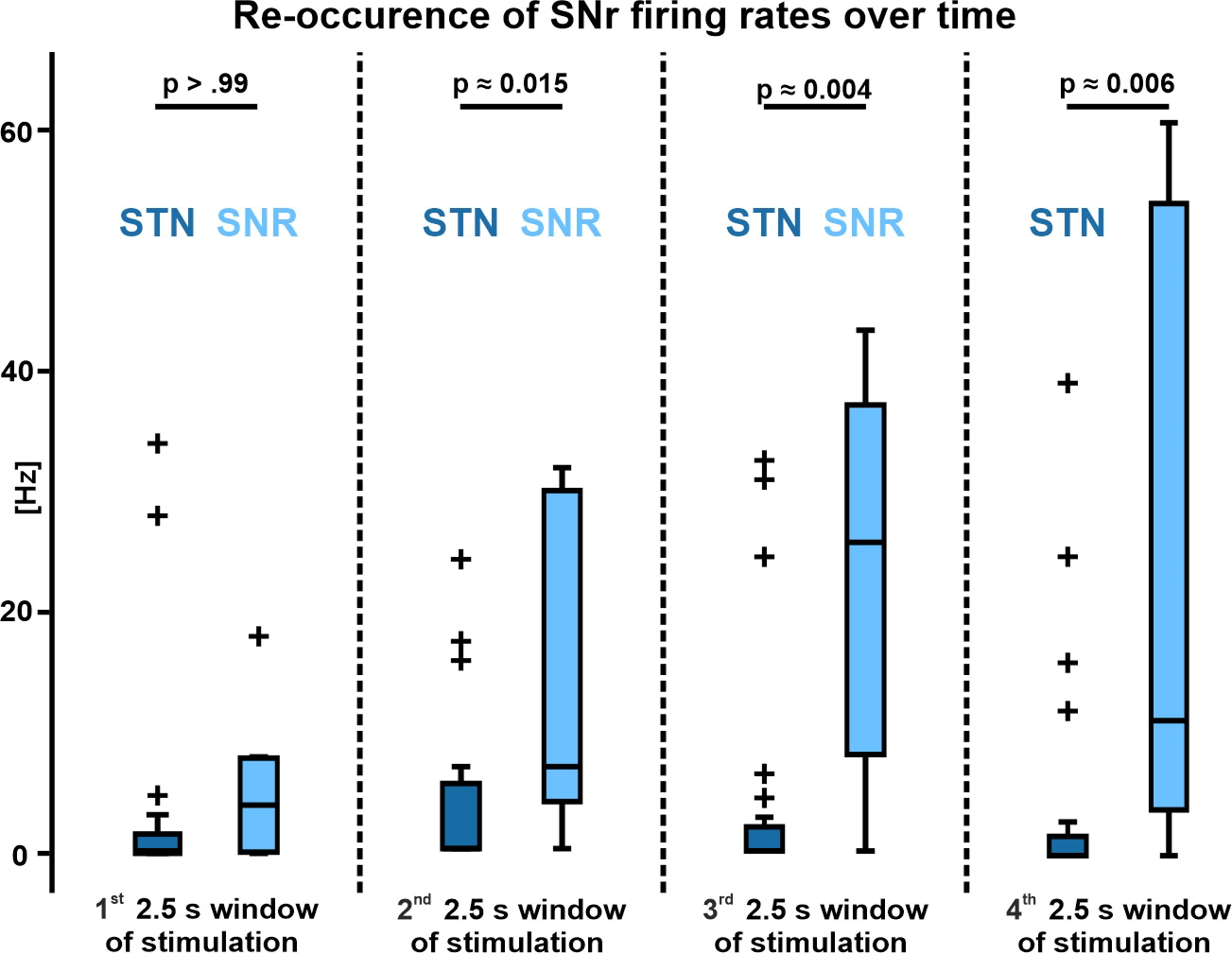

***Supplementary Figure S3: Re-occurrence of SNr firing rates over time.*** Group data show distribution of the return of action potential firing in the SNr as reported on in this results section. The 10 s of stimulation were divided into four 2.5 s windows and data for each neuron is displayed. While there was no difference for the first 2.5 s, suppression of STN neurons was stronger for the other three 2.5 s windows. P-values are taken from serial Bonferroni-corrected (four comparisons) t-tests.
